# A general and powerful stage-wise testing procedure for differential expression and differential transcript usage

**DOI:** 10.1101/109082

**Authors:** Koen Van den Berge, Charlotte Soneson, Mark D. Robinson, Lieven Clement

## Abstract

**Background:** Reductions in sequencing cost and innovations in expression quantification have prompted an emergence of RNA-seq studies with complex designs and data analysis at transcript resolution. These applications involve multiple hypotheses per gene, leading to challenging multiple testing problems. Conventional approaches provide separate top-lists for every contrast and false discovery rate (FDR) control at individual hypothesis level. Hence, they fail to establish proper gene-level error control, which compromises downstream validation experiments. Tests that aggregate individual hypotheses are more powerful and provide gene-level FDR control, but in the RNA-seq literature no methods are available for post-hoc analysis of individual hypotheses.

**Results:** We introduce a two-stage procedure that leverages the increased power of aggregated hypothesis tests while maintaining high biological resolution by post-hoc analysis of genes passing the screening hypothesis. Our method is evaluated on simulated and real RNA-seq experiments. It provides gene-level FDR control in studies with complex designs while boosting power for interaction effects without compromising the discovery of main effects. In a differential transcript usage/expression context, stage-wise testing gains power by aggregating hypotheses at the gene level, while providing transcript-level assessment of genes passing the screening stage. Finally, a prostate cancer case study highlights the relevance of combining gene with transcript level results.

**Conclusion:** Stage-wise testing is a general paradigm that can be adopted whenever individual hypotheses can be aggregated. In our context, it achieves an optimal middle ground between biological resolution and statistical power while providing gene-level FDR control, which is beneficial for downstream biological interpretation and validation.

## Introduction

Next generation sequencing (NGS) technology has become the dominant platform for transcriptome profiling. It is agnostic of genomic annotation, has a broad dynamic range and allows data aggregation on different biological levels (basepair, exon, gene) [1, 2, 3, 4]. Recent developments in read alignment provide fast transcript-level quantification [3, 5, 6] opening the way to assess differential transcript expression (DTE) and differential transcript usage (DTU), which for instance has been shown to be associated with Parkinson’s Disease [7] and resistance to prostate cancer treatment [8]. The dramatic sequencing cost reduction has also enabled researchers to set up studies with complex experimental designs involving many samples [9].

Analysis of DTU, DTE or traditional RNA-seq studies with complex designs typically involve multiple hypotheses of interest for each gene, e.g. for each transcript in a DTU and DTE context or for every treatment effect at each time point and the treatment-time interactions in time course differential gene expression (DGE) studies. The current consensus is to control the false discovery rate (FDR) on the hypothesis level, which we argue to be suboptimal with respect to statistical power and the downstream biological interpretation and validation that typically occurs on a gene level. Soneson et al. [10] has shown that DTE analysis has higher performance when evidence on all individual transcripts is aggregated at the gene level due to the different null hypothesis and the larger amount of data that is available than for tests at the individual hypothesis level. This also occurs for DTU (see Figure 1). Inference using p-values of a DEXSeq [2] analysis on transcript counts also has a lower power than aggregating transcript-level p-values to the gene level prior to FDR calculation. But, the latter does not provide identification of the specific transcripts that are differentially used, thus higher sensitivity comes at the cost of a lower biological resolution.

**Figure 1:**
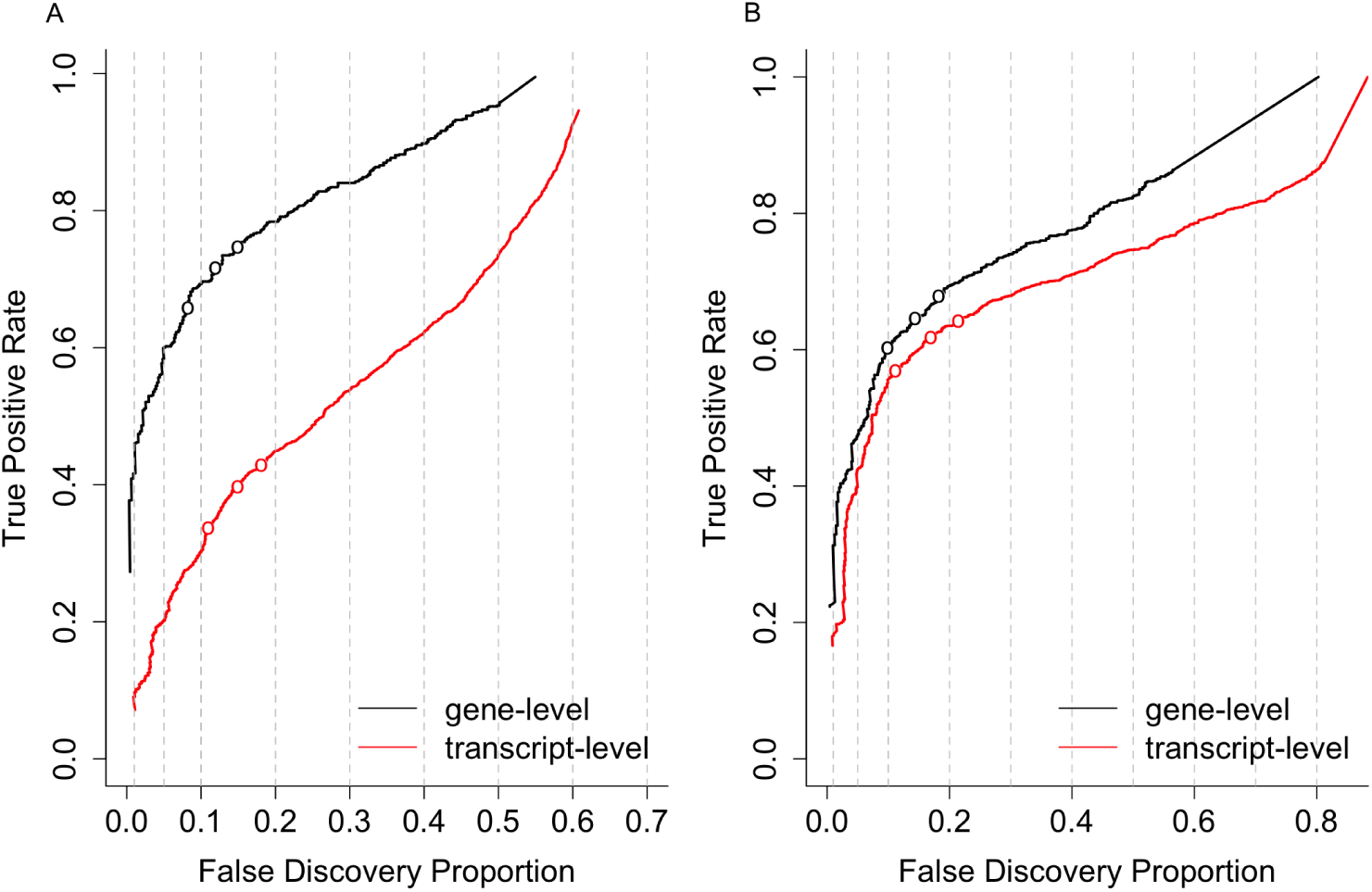
Performance curves for DTU analysis based on two simulation studies. The false discovery proportion (FDP, x-axis) is the fraction of false positive hypotheses over all rejected hypotheses. The true positive rate (TPR, y-axis) represents the fraction of false null hypotheses that have indeed been rejected. The three points on each curve represent working points on a nominal 1%, 5% and 10% FDR. The left panel (A) shows the results from a simulation based on the human transcriptome used in Soneson et al. (2016a) [10] and clearly shows the increased sensitivity for tests that aggregate all transcript hypotheses on a gene level (black curve) in comparison to transcript-level tests (red curve). The right panel (B) shows the results of a simulation study based on *Drosophila melanogaster* performed in Soneson et al. (2016b) [11]. Again, aggregated hypothesis tests have a higher performance, however the difference is smaller in comparison to human, possibly due to the lower complexity of the transcriptome and thus a lower expected number of transcripts per gene for *Drosophila*.

In differential expression (DE) studies with complex designs it is common practice to adopt multiple testing at the hypothesis level. This results in a low power for discovering interaction effects since their standard error is typically much larger than for the main effects. Testing the treatment-time interaction effect in the cross-sectional time series RNA-seq study from Hammer et al., 2010 [12] with limma-voom [13], for instance, returns no significant genes at a 5% FDR level, while more than 6000 genes are flagged when testing for treatment effects within a particular timepoint. Hence, the higher resolution on the hypothesis level comes at the expense of a low power for the interaction effect. In addition, FDR control on the hypothesis level does not guarantee FDR control on the gene level because multiple hypotheses are assessed per gene and the expected ratio of the number of false positive genes to all positive genes in the union across hypotheses will be larger than the target FDR. This can lead to lower success rates of subsequent validation, since many genes without true treatment effects may be considered significant. In the RNA-seq literature, however, there is no consensus on how to combine the enhanced power of aggregation with an adequate resolution for the biological problem at hand. We argue that the multiple hypotheses at the gene level can be exploited in a two-stage testing procedure (Figure 2) [14, 15, 16]. In the screening stage, genes with effects of interest are prioritized using an omnibus test, e.g. a global F-test, a global likelihood ratio test or by aggregating p-values. Assessing the aggregated null hypothesis has the advantages of (1) high sensitivity in a DTU/DTE context, (2) enriching for genes with significant interaction effects in complex DE studies, thereby boosting power; and (3) providing gene-level FDR control. In the confirmation stage, individual hypotheses are assessed for genes that pass the screening stage. Hence, it has the merit to combine the high power of aggregated hypothesis tests in stage I with the high resolution of individual hypothesis testing in stage II.

**Figure 2:**
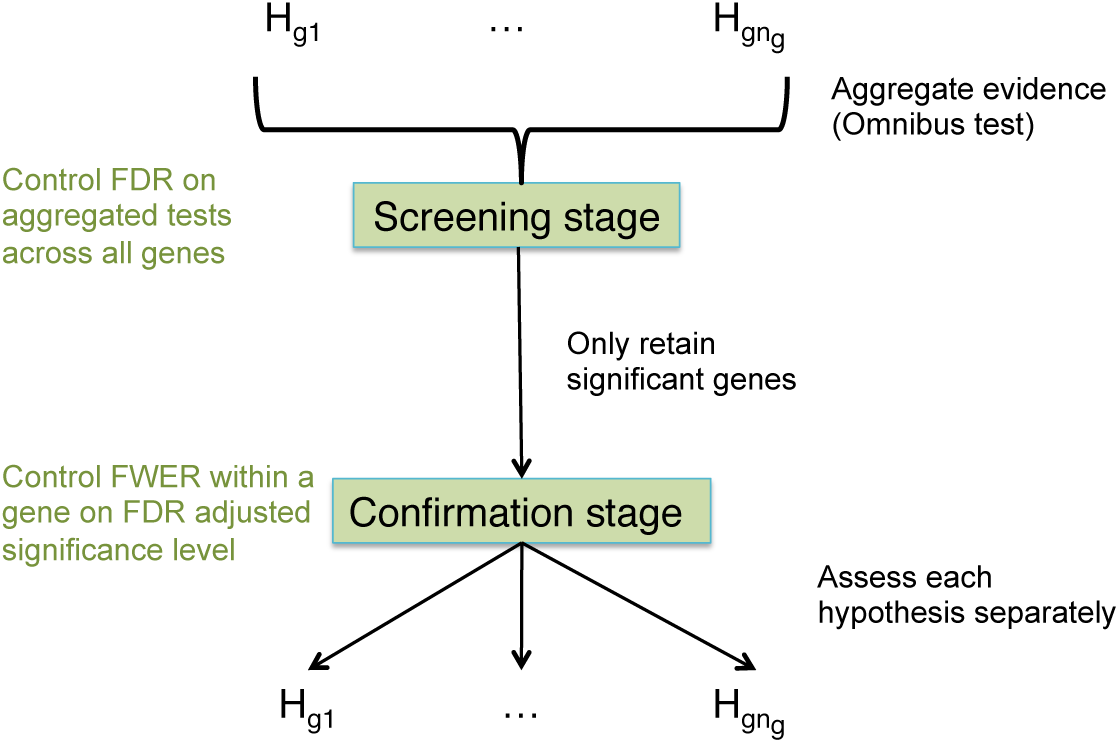
The stage-wise testing paradigm. *n_g_* hypotheses are of interest for gene *g*. In the screening stage, evidence across the hypotheses is aggregated and an omnibus test is performed that controls the FDR across all genes. Genes significant in the screening stage proceed to the confirmation stage, where every hypothesis is assessed separately and FWER is controlled at the adjusted significance level from the screening stage.

The suggested strategy positions itself in the larger framework of stage-wise testing procedures for high-throughput experiments. Lu et al. [17] previously proposed a two-stage strategy for microarrays based on mixed models, which is inapplicable to NGS data due to the violation of the distributional assumptions. Jiang & Doerge [14] proposed a generic two-stage differential expression analysis procedure where the first stage corresponds to testing a global null hypothesis, after which post hoc contrasts are tested only for the significant genes. The algorithm, however, requires either parametric assumptions or computer intensive resampling-based techniques. Heller et al. [15] proposed a two-stage procedure in the context of gene set enrichment analysis (GSEA) for microarray data: in the screening stage, the global null hypothesis is tested for each gene set and the procedure then proceeds to test for differential expression of individual genes within discovered gene sets. It was shown that their procedure controls the overall FDR (OFDR) [18] under independence, positive regression dependency and dependencies that are typically occurring in microarrays [19, 20, 15]. They define the OFDR as the expected proportion of falsely discovered gene sets over all discovered gene sets and a falsely discovered gene set as a gene set for which at least one null hypothesis (including the screening hypothesis) is falsely rejected [18, 15].

In this contribution, we port the ideas developed in Heller et al. [15] to the DTU, DTE and DE problem in NGS experiments with simple and complex designs by (1) replacing “a gene set” in their procedure by “a gene”,(2) aggregating evidence across all individual hypotheses per gene in the screening stage and (3) assessing each individual hypothesis on the discovered genes. In our context, the OFDR thus controls the FDR at the gene level and we argue this to be the most appropriate error rate in complex high throughput experiments due to its link with subsequent gene-level interpretation and validation by the biologists. We further improve the power in the second stage of the Heller method by developing multiple testing procedures specifically tailored to the problem at hand. Similar to Meijer & Goeman [21, 22], our methods exploit the logical relations between the hypotheses that have to be assessed within each gene to reduce the multiple testing burden in the second stage. The procedure is powerful, easy to implement and we will show that it provides an optimal middle ground between statistical power and resolution on the biological research questions for both DGE, DTE and DTU analyses. The method has been implemented in an R package `stageR` available at https://github.com/statOmics/stageR.

## Results

We evaluate the stage-wise testing procedure in DGE, DTE and DTU applications on both synthetic and real data. The results of the two-stage method are compared to the current consensus of a data analysis workflow in the specific applications. First, we verify the gene-level FDR control and power for complex DGE experiments on simulated data upon which we confirm the simulation results on real data. For DTE and DTU analysis, we show how the stage-wise testing procedure maintains high sensitivity on the gene level while simultaneously maintaining a high resolution on the biological results. Additionally, we show that performance on the transcript level is at least as good as with a regular transcript-level analysis. By analysing a real prostate cancer dataset, we illustrate how the combination of gene and transcript-level results provides a rich resource for follow-up biological interpretation and validation.

## Differential gene expression

### Simulation study

The simulation study is set up according to the Hammer study [12], a full factorial design with a factor time (timepoint 1, timepoint 2) and treatment (control, spinal nerve ligation (SNL)) with two levels each. RNA-seq counts for 13,000 genes are simulated, 2,000 genes have a constant fold change between treatment groups over time (main effect contrast), 2,000 genes show DE in only one timepoint (1,000 genes for every timepoint) and 1,000 genes are differentially expressed in both timepoints with a different fold change between the timepoints(interaction effect). 30 datasets are simulated with either five or three biological replicates in every treatment x time combination. The hypotheses of interest are DE at timepoint 1 and/or timepoint 2 (5,000 genes) and testing for a change in DE between timepoint 1 and 2 (3,000 genes with a real treatment x time interaction). A conventional approach assesses each of these hypotheses separately. Our two-stage approach, however, considers a test with an aggregated null hypothesis in the screening stage, i.e. that there is no effect of the treatment whatsoever. The individual hypotheses are only assessed in the confirmation stage for genes that passed the screening stage,i.e. for genes showing evidence for a treatment effect. We model the read counts by (generalized) linear models with a treatment effect, time effect and treatment x time interaction using the limma-voom (edgeR) [13, 23] framework and compare the conventional approach to a two-stage analysis in terms of FDR control, OFDR control and power.

Figure 3 shows the results for the limma-voom analysis of the simulated datasets with five biological replicates. Both the standard and stage-wise analysis control the FDR on the hypothesis level. This is expected for the standard approach, but is generally not guaranteed for the stage-wise approach since the latter is designed to control the OFDR. Both procedures reject approximately the same number of individual hypotheses for the treatment effects within each timepoint. The standard analysis, however, discovers a higher number of genes with only one significant contrast of interest (Additional File 1: Figures S4, S9), indicating that the stage-wise approach enriches for genes with multiple effects. Moreover, the standard analysis leads to poor OFDR control, while the stage-wise analysis controls the OFDR at its nominal level, indicating the superiority of a stage-wise testing approach to prioritise candidate genes for further analysis. Biological validation and interpretation using tools like gene set enrichment analysis often occurs at the gene level, which motivated us to compare the fraction of null genes (i.e. genes where all null hypotheses are true) in both candidate gene-lists. When decomposing the OFDR false positive genes in null genes and genes with a treatment effect but where one of the true null hypotheses has been falsely rejected, we observe that the decrease in the fraction of false positives is more pronounced for the stage-wise method than for the conventional analysis (Figure 3). Thus, in addition to providing gene-level FDR control, the fraction of null genes among the false positive list is also lower for the stage-wise method, which will eventually limit resources wasted towards the validation of false positive genes in follow-up experiments. The results of the negative binomial edgeR analysis are qualitatively similar to the limma-voom analysis with the exception that FDR control can be rather liberal for edgeR, especially for small sample sizes (Additional File 1: Figures S1, S6).

**Figure 3:**
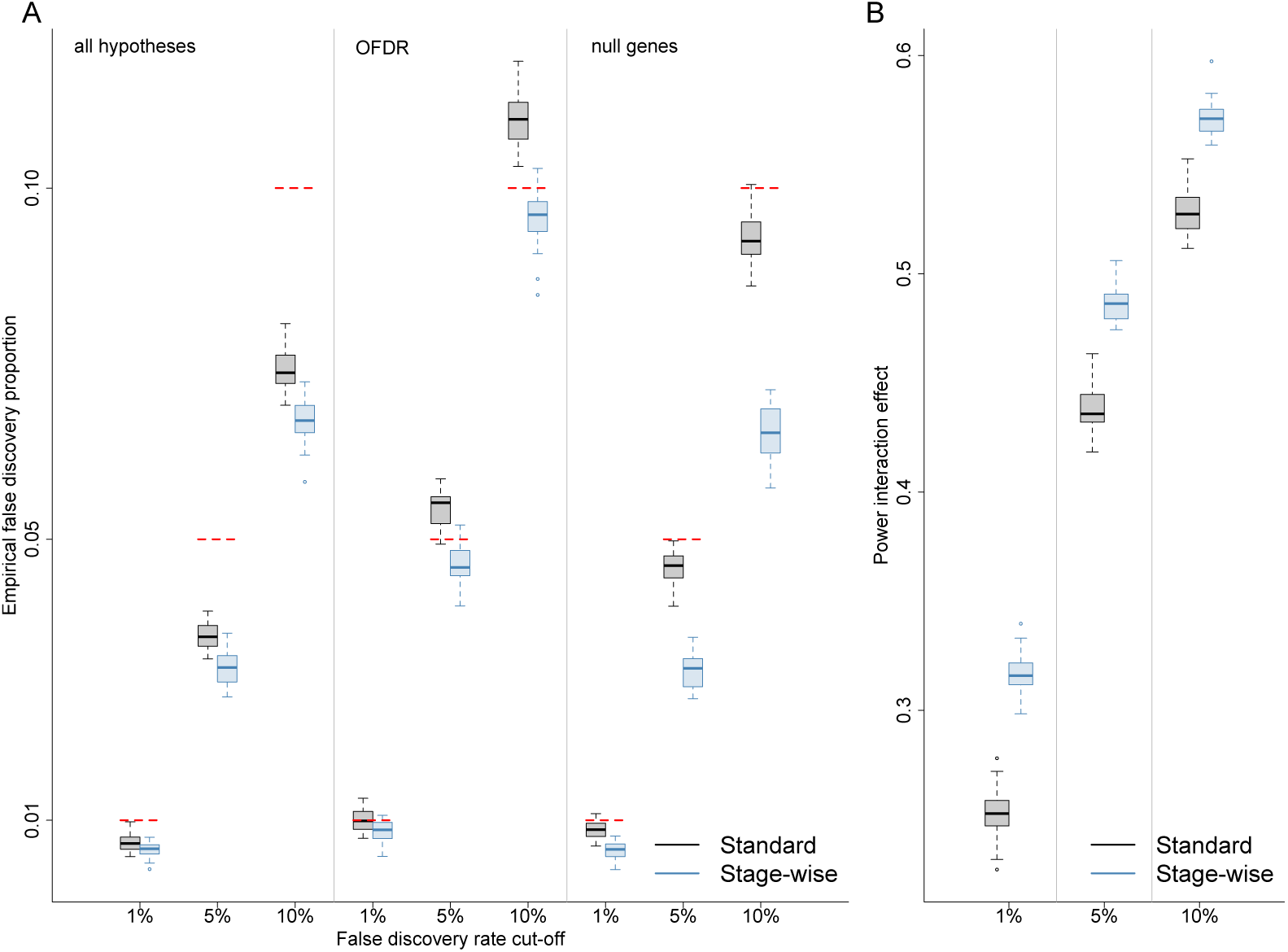
DGE simulation study results for the limma-voom analysis with five replicates in every treatment x time combination. (A) FDR and OFDR control for the stage-wise (blue) and conventional (black) methods. The false discovery proportion (FDP) is assessed in 30 simulations which allows us to evaluate the FDR as the mean over all FDPs. Both the standard and stage-wise approach control the FDR over all hypotheses, which is expected for the conventional method but not for the stage-wise analysis. OFDR is controlled in the stage-wise analysis, while a standard hypothesis-level approach is too liberal on the gene level. In addition, the fraction of null genes among the OFDR false positive list is lower for the stage-wise testing procedure, which shows that it is advantageous in terms of e cient biological validation of the results. (B) Power evaluation for the treatment × time interaction effect. The stage-wise method boosts power for the interaction effect through the enrichment of interaction genes in the screening stage.

In the introduction, we illustrated the low power of the conventional method for discovering interaction effects in RNA-seq experiments, and we observe a similar behaviour in the simulation study. Our two-stage method, however, enriches for genes with interaction effects by aggregating evidence across hypotheses in the screening step. This results in a power boost of 5% in the simulation study with ve replicates (Figure 3). This power increase is even more pronounced if a lower number of replicates are available (Additional File 1: Figures S5, S6). Note, that the stage-wise and standard analysis show equivalent performance for the treatment effects within each time point (Additional File 1: Figures S2, S3, S7, S8). Hence the stage-wise analysis (1) returns a lower number of false positive genes and false positive null genes, (2) provides a higher power for interaction effects while (3) maintaining the same performance for the main effects.

### Case study

We also re-analyse the Hammer dataset [12] with the two-stage method using limma-voom. The results are very similar to the DGE simulation study, suggesting good quality of the simulated data: contrasts involving main effects have a similar number of significant genes between standard and stage-wise procedures while the latter again finds many more significant genes when testing for the interaction effect (Table 1). While the conventional analysis did not find any genes when testing for the interaction effect, the stage-wise method retrieves 665 significant genes. The results of negative binomial count regression with edgeR are in line with the limma-voom analysis (Additional File 1: Table S1). Note, however, that edgeR does find 51 significant genes for the interaction test in a conventional analysis, which however may be a result of the higher power associated with edgeR’s rather liberal FDR control for experiments with small sample sizes (Additional File 1: Figure S6).

85 genes had only passed the screening stage while none of them had a significant effect in the confirmation stage or by the standard analysis. Their expression profile reveals a moderate fold change with respect to the treatment that remains stable over time (Additional File 1: Figures S10, S11), again indicating the higher sensitivity of the overall test in the screening stage. All genes, however, could be retrieved when testing for a contrast that quantifies the average fold change (i.e. average DE between SNL and control over time) confirming the biological relevance of the stage-wise testing approach, even for the genes without significant effects in the second stage. Note, that incorporating the test for the average fold change does not alter the FWER correction of the Shaffer method (see Methods) and can be adopted in the confirmation stage without compromising the power on the other contrasts of interest.

**Table 1:**
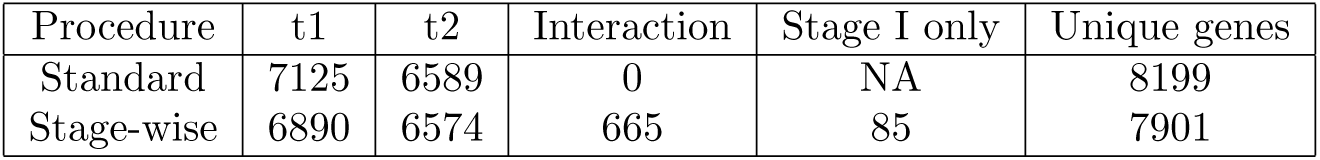
Number of genes found in the Hammer dataset on a 5% FDR level in the limma-voom analysis. 12,893 genes were considered in the analysis.

| Procedure | t1 | t2 | Interaction | Stage I only | Unique genes |
| --- | --- | --- | --- | --- | --- |
| Standard | 7125 | 6589 | 0 | NA | 8199 |
| Stage-wise | 6890 | 6574 | 665 | 85 | 7901 |

## Differential transcript usage and differential transcript expression

### Simulation study

We adopt the *Drosophila melanogaster* and *Homo sapiens* simulation studies from Soneson et al. [11] for both DTE and DTU analysis. In the original simulation study DTU was simulated by flipping the proportions of the two most abundant transcripts, while in real data we often observe more than two significant transcripts per gene (e.g. Figure 5). Hence, we have extended the simulation study to accommodate alternative splicing patterns across multiple transcripts per gene and we also allow low-expressed genes to be simulated as differentially expressed or used. We compare the performance of the standard and stage-wise procedures both on the transcript and gene level by performing transcript-level tests or aggregating transcript-level p-values, respectively. We did not consider analyses based on gene-level aggregated counts in the screening stage because this approach would fail to find DTU for genes with constant output between conditions. The gene-level test is superior to the transcript-level test in terms of sensitivity for both DTE (Figure 4) and DTU (Additional File 1: Figure S12) which motivates the stage-wise testing procedure. Furthermore, by leveraging the power in the gene-level test to the transcript-level analysis in the confirmation stage, the stage-wise analysis also has higher performance on the transcript level in a DTE analysis (Figure 4) and is at least on par in a DTU analysis (Additional File 1: Figure S12) compared to a regular transcript-level approach. The stage-wise transcript-level tests do not only result in increased performance but additionally provide a better FDR control on the transcript level. Due to the better FDR control, the stage-wise analysis will not necessarily find more transcripts as compared to a transcript-level analysis, but the number of true positive transcripts for a fixed fraction of false positive transcripts in the rejected set should be at least identical or higher in the stage-wise analysis. We confirm the observation made in Soneson et al. [11] that FDR control deteriorates severely in human as compared to fruit fly and support their hypothesis that this is related to transcriptome complexity.

**Figure 4:**
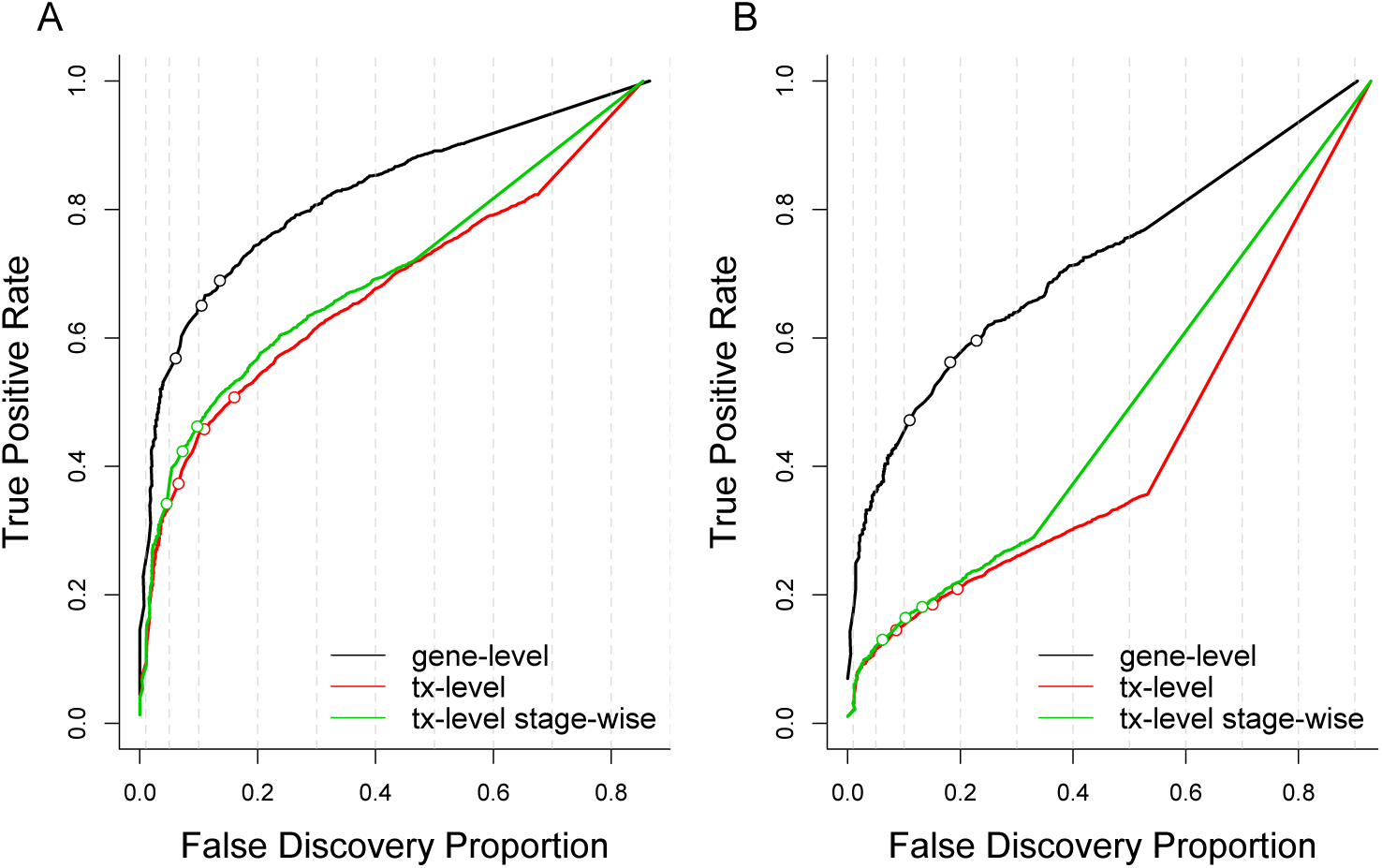
FDP-TPR performance curves for DTE analysis of the simulated data. Red curves represent transcript-level tests and black curves represent tests based on p-values aggregated at the gene level. The green curve represents the stage-wise transcript-level analysis. The three open circles on the curves represent working points for a target FDR of 1%, 5% and 10%. (A) Performance curve for the *Drosophila* simulation shows an inflated FDR for both the aggregated analysis on the gene level and a transcript-level analysis. The stage-wise transcript-level analysis has increased performance and additionally provides a better FDR control. (B) Performance curve for the human simulation shows even worse FDR control on all levels, which is in line with previous publications [11]. Similar to the *Drosophila* simulation, the transcript-level stage-wise analysis shows somewhat higher performance and provides a better FDR control.

**Figure 5:**
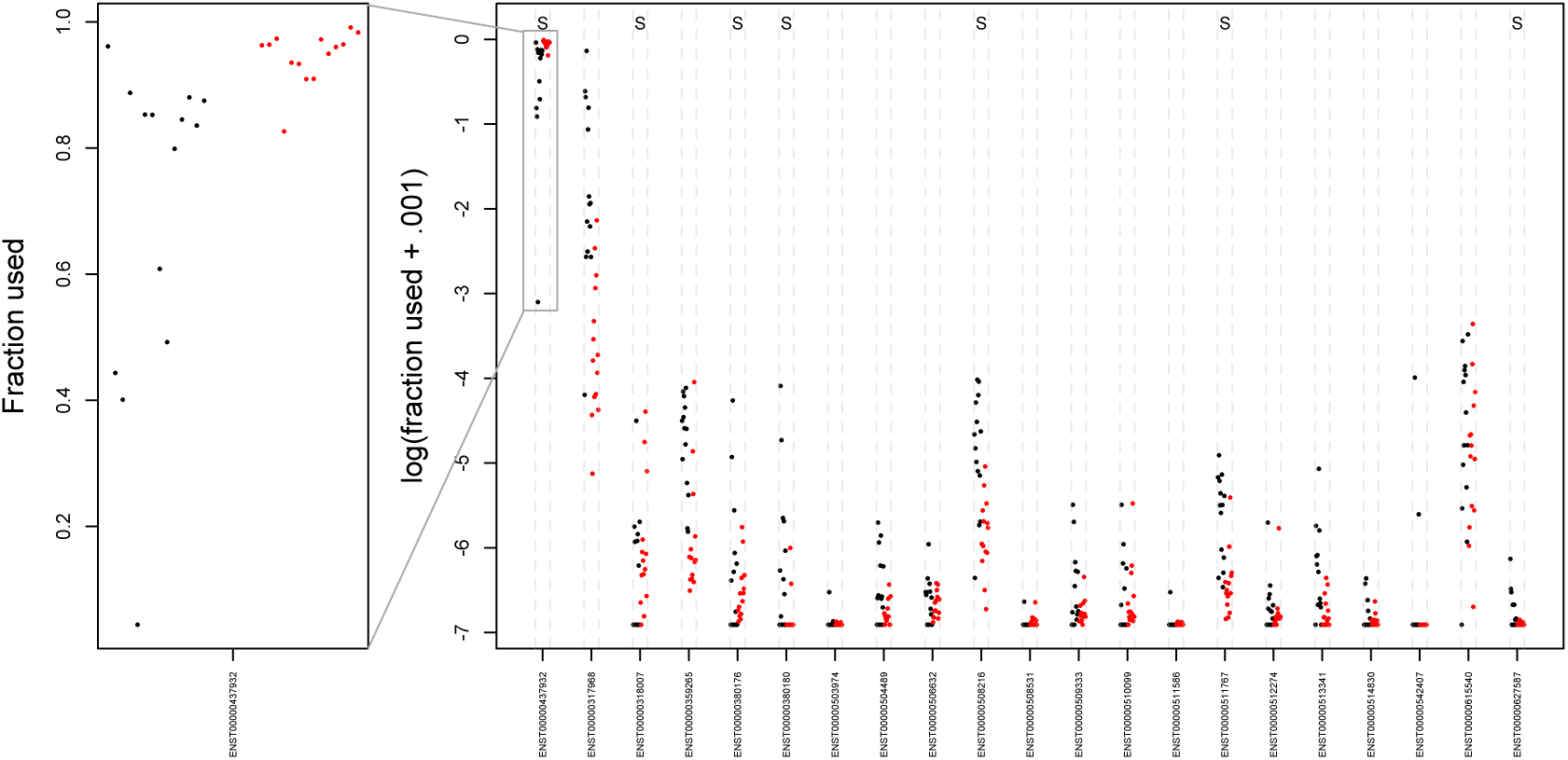
Expression pattern of the PDLIM5 gene in the case study. The used fraction for every transcript is relative to the total expression of the genomic locus for a respective sample. Black points represent normal tissue and red points represent tumoral tissue. The left panel (original scale) shows the dominant transcript that is additionally upregulated in tumoral tissue. The right panel shows the usage pattern on the log scale for all transcripts and shows that the upregulation of the dominant transcript is compensated for by a downregulation of multiple other transcripts. significantly differentially used transcripts according to the stage-wise analysis are indicated with an S at the top of the plot.

### Case study

We analysed a prostate cancer dataset [24] from 14 Chinese tumour-normal matched samples for DTU and DTE. We use DEXSeq [2] for assessing DTU by providing transcript-level expression estimates instead of exon bin quantifications hence we fit transcript-level negative binomial models. In the screening stage, inference is based on transcript-level p-values aggregated to the gene level and transcripts of significant genes are confirmed using individual transcript-level tests. In the DTU analysis, we filter genes with only one transcript, which leaves 18,479 genes with a median of 6 transcripts per gene. On a 5%target OFDR level, the stage-wise testing analysis finds 4,752 significant genes in the screening stage and confirms 6,772 significant transcripts in the confirmation stage. Similarly, DTE is assessed using transcript-level negative binomial models implemented in edgeR [25] and gene-level tests are performed by aggregating transcript-level p-values. For DTE, no ltering is required so we consider 32,499 genes with a median of 2 transcripts per gene, although the analysis may bene t from independent low abundance ltering [26]. The stage-wise analysis finds 4,842 significant genes in the screening stage and confirms 5,236 transcripts in the second stage on a 5% target OFDR level.

The literature concerning alternative splicing has often focussed on isoform dominance, where it is hypothesised that many genes have dominant isoforms and a major mechanism of differential transcript usage would be a switch of dominance between two transcripts [27]. We find that two out of five genes (CYP3A5 and LPIN1) with a gene-wise q-value equal to zero have previously been associated with prostate cancer [28, 29] and indeed both genes seem to correspond to a switch in the dominant isoforms (Additional File 1: Figure S13). However, for both genes the association with prostate cancer was based on differential expression analysis while we show that instead the underlying pattern of the previous results is due to differential transcript usage. We also observe that complex genes with many transcripts are much more likely to be flagged in a DTU analysis as compared to genes with a low number of isoforms (Additional File 1: Figure S15). 10% of significant genes with at least three transcripts have three or more transcripts confirmed as differentially used in the confirmation stage, providing accumulating evidence of more complex biological splicing patterns. According to the stage-wise testing method, two genes have seven or more differentially used transcripts on a 5% target OFDR level. The transcript usage pattern for one of them, the PDLIM5 gene, corresponds to an upregulation for a dominant transcript that is compensated for by a downregulation of multiple others (Figure 5, Additional File 1: Figure S14). Remarkably, a SNP (rs17021918) in this gene was also found to be associated with prostate cancer in previous studies [30, 31], among which a large-scale multi-stage GWAS study that provided robust associations across multiple populations [30]. When compared to a log-additive model, which is the most common model for association of SNPs with disease and assumes an additive effect of the log-odds on disease for each copy of the allele, the effect of rs17021918 exceptionally showed no difference in risk on prostate cancer between heterozygotes and homozygotes. Furthermore, the SNP lies in the intronic region of the PDLIM5 gene and could contribute to alternative splicing patterns observed in prostate cancer cells as compared to normal cells instead of providing allele dosage effects on prostate cancer risk.

## Discussion

We adapted the two-stage procedure of Heller [15] as a general inference paradigm to provide powerful statistical testing and FDR control for problems that allow hypotheses to be aggregated. We tailored it towards modern RNA-seq applications assessing many hypotheses per gene and showed that it is superior in terms of interpretation, sensitivity and specificity. The screening stage considers an omnibus test aggregating evidence across all hypotheses for every gene. This boosts the sensitivity for effects that have a relative low power, e.g. interactions in studies with complex designs and for picking up genes with differential transcript expression and transcript usage. Note, that screening also results in a shift of the type of genes that are returned. The omnibus test might dilute the evidence of genes with a moderate effect for a single contrast (transcript) by aggregating it with the true null hypotheses for the remaining contrasts (transcripts). The loss of these genes, however, is compensated by the discovery of additional genes with moderate effect sizes for multiple contrasts of interest (transcripts) in studies with complex designs (DTE/DTU applications), e.g. the genes picked up in the Hammer study with a stable and moderate differential expression over time in rats with spinal nerve ligation compared to controls. Upon screening, individual effects/transcripts are further explored for the discovered genes. For RNA-seq applications, we also optimized the power of the confirmation stage by accounting for logical relations between the hypotheses that have to be assessed within each gene.

We have focussed on time-series data in the application of the two-stage method on DGE analysis. However, the two-stage procedure is generic and can be applied to any design. For example, a DGE study that compares three drugs (e.g. a new drug, the current state of the art and a placebo) would require exactly the same data analysis paradigm as the Hammer dataset: three different hypotheses of interest (mean differential expression between the drugs) and according to Shaffer’s MSRB procedure no correction is needed in stage II for FWER control. The extension to more complex designs is trivial when Holm’s method is used in stage II. However, a good understanding of the logical relations among the hypotheses is required for implementing Shaffer’s MSRB procedure so as to obtain maximal power.

The standard transcript-level and gene-level approaches for DTU/DTE analysis show an inflated FDR in all simulations. The stage-wise transcript-level analysis provides better FDR control although it is often still inflated, especially for the human simulations. Indeed, the distribution of human transcript-level p-values is non-uniform for the higher p-values (Additional File 1: Figure S16) suggesting invalid statistical inference, while a proper p-value distribution is observed for the *Drosophila* simulation (Additional File 1: Figure S16). Soneson et al. [11] suggested that the inflated FDR is related to transcriptome complexity and have shown that FDR control can become problematic for genes with many transcripts, which is partly due to the uncertainty associated with read attribution to similar transcripts. In addition, recent work has shown that transcript abundance estimates are infested with systematic errors as a result of a failure to model fragment GC content bias [32], leading to false positive transcript abundance results. Despite the computational advantage of light-weight algorithms like Salmon and kallisto [6, 3], many problems remain for correct estimation of transcript abundances, all of which may contribute to inflated FDR in transcript-level analyses. Even if the true transcript abundances could be obtained, the statistical inference engine of DEXSeq relies on large sample assumptions that are often not met in reality and further method development is required to provide correct FDR control in a DTE/DTU context. Note, however, that our method is very general and can be easily adopted as new frameworks for DTE and DTU become available.

The DTU case study highlights the biological relevance of differential transcript usage in oncology research. It shows that prioritising genes in a first stage and subsequently confirming transcripts for the significant genes provides an elegant approach to DTU analysis, upon which biological interpretation of the results may follow. We confirm that a switch between dominant isoforms is a common pattern in differential transcript usage, but additionally provide evidence that the field may benefit from considering more complex splicing mechanisms as was shown for the PDLIM5 gene. The stage-wise testing method provides an optimal data analysis strategy for discovering genes with dominant isoform switches as well as for genes with more subtle changes in differential transcripts.

It is also important to stress that the stage-wise testing procedure has the merit to control the FDR at a gene level, which we claim to be beneficial over FDR control at a hypothesis level in experiments involving many hypotheses per gene. The gene is the natural level for downstream analysis, e.g. gene set enrichment analyses and subsequent biological validation experiments. These might be compromised when using traditional hypothesis-level FDR control since the union of all genes found across hypotheses tends to be enriched for false positive genes for which all null hypotheses are true and that are not of interest to the biologist.

The Heller method controls the expected fraction of rejected genes with at least one false positive hypothesis and uses a FWER correction within a gene in the confirmation stage. If many hypotheses are of interest for all genes then a less stringent error measure can be used as proposed by Benjamini & Bogomolov [16], e.g. by using FDR control across the hypotheses within a gene in the confirmation stage. Their method has similarly been extended and applied to a large-scale multi-trait genome-wide association study [33]. However, one then loses the interpretation of a gene-level FDR control, which is very useful in our context.

In this contribution we have deliberately chosen a stage-wise approach with nested hypothesis tests in the screening and confirmation stage. The method of Heller, however, does not imply the use of an omnibus test in the screening stage. In a DTE context, for instance, we also might opt to aggregate the transcript-level counts at a gene level instead of aggregating evidence over transcript-level hypotheses. We, however, feel that this will obscure the interpretation. In the latter approach, it is unclear how the screening step will enrich for the hypotheses of interest at a transcript level. For instance, genes with DTU and equal overall expression can exhibit a very clear DTE signal but are bound to fly under the radar when aggregating counts. The null hypothesis of the omnibus test in the screening stage has a natural interpretation that none of the effects of interest occur for a particular gene, versus the alternative that at least one effect is present. Hence, the screening stage will enrich for genes with effects that will be further explored in the confirmation stage.

## Conclusions

We have introduced two-stage testing as a general paradigm for assessing high throughput experiments involving multiple hypotheses that can be aggregated, which is implemented in the R package stageR (https://github.com/statOmics/stageR). We optimized the procedure towards RNA-seq applications: differential transcript expression, differential transcript usage and differential gene expression analysis with simple and complex experimental designs. We have shown that the procedure controls the overall FDR, which we argue to be the natural error rate in high-throughput studies: in our context the OFDR gets the interpretation of a gene-level FDR and shares a close link with the subsequent biological validation experiments and interpretation of the results, e.g. gene set enrichment analyses. The omnibus test in the first stage boosts the power when testing for interaction effects in DE studies with complex designs, without compromising power on the remaining contrasts. For DTU and DTE analysis,the two-stage method gains from the high performance of FDR control upon aggregating transcript-level p-values to the gene level. Specific transcripts can be identified in the subsequent confirmation stage for genes passing the screening stage. Hence, the two-stage procedure naturally unites the highest level of resolution on the biological problem with the superior power of aggregated hypothesis tests. In addition, the two-stage transcript-level analysis is on par(DTU) or has higher performances (DTE) than a conventional transcript-level analysis while providing better FDR control. We have used the two-stage testing procedure to prioritise interesting genes in a case study on prostate cancer and we illustrated the potential of DTU analyses in the context of cancer research.

## Methods

**Two-stage testing procedure**

The two-stage testing procedure that is proposed in this contribution was introduced by [15] for assessing gene set enrichment analysis. We adapt the procedure and formulate it more general. Suppose we have a dataset that consists of *G* genes. For every gene *g*, we are interested in testing *n_g_* null hypotheses *H_1g_*,…,*H_n__g_g*. In the screening stage the global null hypothesis 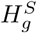 is assessed, i.e. that all *H_1g_*,…,*H_n_g_g_* are true against the alternative hypothesis that at least one hypothesis *H_ig_* is false. The confirmation stage consists of assessing all individual hypotheses *H_1g_*,…,*H_n_g_g_* for each gene that passed the screening stage. The procedure proceeds as follows:

### 1. Screening Stage

- Assess the screening hypothesis 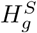 for all genes in the set *G*.
- Let 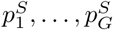 be the unadjusted p-values from the screening stage test.
- Apply the Benjamini Hochberg (BH) FDR procedure [34] to 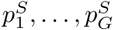 at FDR level α*_I_*. Let R be the number of rejected screening hypotheses.

### 2. Confirmation Stage

For all *R* genes that pass the screening stage.

- Let *α_II_* = *Rα_I_* / *G* the BH-adjusted significance level from the first stage.
- Let *p_1g_*,…,*p_n_g_g_* be the p-values from *H_1g_*,…,*H_n_g_g_* for gene *g*.
- Adopt a multiple testing procedure to assess all *n_g_* hypotheses while controlling the within gene family wise error rate (FWER) at the adjusted level *α_II_*.

Heller et al. [15]prove that the procedure controls the OFDR under independence between genes, an assumption that is required for the BH procedure. The BH procedure has been proven to be also valid under positive regression dependency [19] and it has additionally been shown to be valid under typical microarray dependencies between the genes [20, 35].

Note, that any FWER correction procedure can be used in stage II. We, however, propose the use of the Shaffer modified sequentially rejective Bonferroni (MSRB) method [36] when logical relationships amongst the hypotheses exist. The MSRB method is a modi ed Bonferroni procedure that accounts for the logical relationships among the hypotheses. Like the regular Bonferroni procedure, it is highly flexible and easily used in nonstandard situations of dependency [36]. In brief, the procedure works as follows. Suppose we have filtered the genes that pass the screening stage. Let *p*_(1)*g*_,…,*p*_(*n_g_*)*g*_ be the sorted unadjusted p-values in the confirmation stage for gene *g* where *p*_(1)*g*_ ≤ *p*_(2)*g*_ ≤… ≤ *p*_(*n_g_*)*g*_. The method works sequentially over the sorted p-values: suppose that the first *j* − 1 hypotheses have been rejected, we then compare *p*_(*j*)*g*_ to *α*_*II*_/*t*(*j*) where *t*(*j*) equals the maximum number of remaining hypotheses that still could be true given that the first *j* − 1 hypotheses are false. *t_j_* is never greater than *n* − *j* + 1 and therefore the MSRB procedure uniformly outperforms the Holm [37] method.

In a standard setting, the first p-value will be compared to *α*_*II*_/*t*(1) = *α*_*II*_/*n_g_*. However, within a two-stage procedure, we know that for every gene in the confirmation stage there is at least one effect, otherwise the screening hypothesis has been falsely rejected. Therefore, the MSRB procedure can be further modified such that the first p-value can be compared to *α*_*II*_/(*n_g_* − 1) hereby boosting power for the most significant test. Below, we show how *t*(*j*) might further reduce according to the speci c context.

## Differential gene expression

### Case study

The Hammer dataset [12] was downloaded from the ReCount [38, 39] project website (http://bowtie-bio.sourceforge.net/recount/). In this experiment, rats were subjected to a spinal nerve ligation (SNL) and transcriptome profiling occurred at two weeks and two months after treatment, for both the SNL group and a control group. Two biological replicates are used for every treatment × time combination. An independent filtering step [26] is performed prior to the analysis after which we retain 12,893 genes with adequate expression (counts per million larger than 2) in at least 2 samples. Data was normalised using TMM normalisation [23] to adjust for variations in sequencing depth and mRNA population. We analyse the data using a log-linear model implemented in limma-voom [13] and hypothesis testing is performed through moderated t-tests and F-tests. Additionally, we re-analyse the data using the negative binomial model implemented in edgeR [25] where hypothesis testing is performed through likelihood ratio tests. We assess (a) the treatment effect at the first timepoint, (b) the treatment effect at the second timepoint and (c) the treatment × time interaction using a contrast for the differential expression at the first and second timepoint and a difference in fold change between the two timepoints, respectively. For the standard analysis, every contrast has been assessed on a 5% target FDR level as was the screening hypothesis in the stage-wise analysis. When a gene passed the screening hypothesis, only one null hypothesis can still be true: there has to be DE at timepoint 1 or timepoint 2; if the DE only occurs on one timepoint there also exist an interaction; if DE occurs at both timepoints, the *H*_0_ of no interaction can still be true. Hence, *t*(3), *t*(2) and *t*(1) = 1 in the Shaffer MSRB method for the Hammer experiment and no additional FWER correction is required in the confirmation stage.

### Simulation study

The simulation study is designed to mimic the Hammer dataset [12]. The raw count table was downloaded from the ReCount project [38, 39] website (http://bowtie-bio.sourceforge.net/recount/). We simulated realistic RNA-Seq data based on the framework provided by [40] with some minor adjustments which allow us to link a gene’s characteristics over different timepoints, unlocking simulation of cross-sectional time-series DGE data. Gene-wise means *μ_g_* and dispersions *ϕ_g_* are estimated from the larger Pickrell dataset [41] for more efficient estimation and are jointly sampled for simulation, respecting the mean-variance relationship of RNA-Seq data. We simulate 13,000 genes (equivalent to the number of genes analysed in the Hammer data) according to a negative binomial model for two timepoints and two conditions. We considered two sample sizes: simulated datasets with either ve or three biological replicates for every treatment × time combination. We simulate 2,000 genes with a constant fold change between control and treatment for both timepoints, 2,000 genes with time-specific differential expression between treatment and control (1,000 genes for every timepoint) and 1,000 genes with a different fold change between timepoints (i.e. significant treatment × time interaction effect). All fold changes were set at 3 or 1/3, balanced in every contrast.

As in the case study, we use limma-voom [13] and edgeR [25] for differential expression analysis. Following [15], we define a false positive gene as a gene where at least one of the null hypotheses (including the screening hypothesis) is falsely rejected and use this criterion to define the OFDR.

## Differential transcript usage and differential transcript expression

### Simulation study

The simulation study is adapted from Soneson et al. [11]. In brief, RSEM [42] generates paired-end sequencing reads with a length of 101 bp based on parameters that are estimated from real RNA-Seq data. The transcripts per million (TPM) expression levels and relative isoform abundances are estimated from real fastq-files with RSEM: as in the original simulation study, we used sample SRR1501444 (http://www.ebi.ac.uk/ena/data/view/SRR1501444) for the Drosophila simulation and sample SRR493366 (http://www.ebi.ac.uk/ena/data/view/SRR493366) for the human simulation. Based on a mean-dispersion relationship derived from two real datasets (Pickrell [41] and Cheung [43] dataset, see [44]), every estimated expression level is matched with a corresponding negative binomial dispersion value for each gene. We scale the TPM values according to the desired library size to derive the gene-wise expected count and simulate counts from a negative binomial distribution. Two conditions were considered in the simulation study and five samples were simulated in each condition. A Dirichlet distribution was used to simulate relative isoform abundances in each sample. We simulate 1,000 genes with differential transcript usage. The genes with differential transcript usage were selected randomly from the subset of genes with expected gene count above 5 and at least two expressed isoforms. The number of differentially used transcripts within the gene is sampled ranging from a minimum of 2 up to a random number drawn from a binomial distribution with size equal to the number of transcripts and succes probability 1/3. We introduce DTU by randomly flipping the proportions between the differentially used transcripts. In addition to the DTU genes we simulate 1,000 DTE genes, where all transcripts from a gene are differentially expressed with fold changes drawn from a truncated exponential distribution. The simulated fastq files were mapped to the *Drosophila melanogaster* (*Homo sapiens*) transcriptome derived from the BDGP5.70 (GRCh37.71) primary genome assembly using kallisto [3].

### Differential transcript usage

Prior to the analysis, we round the estimated isoform-level counts to the closest larger integer and discard genes with only one transcript and transcripts with no expression over all samples. The count matrix was then used as input to DEXSeq [2]. DEXSeq estimates size factors as in DESeq [45] for data normalization. A transcript-wise negative binomial generalized linear model is fitted and changes in relative usage between the conditions are assessed by testing the `transcript: condition` interaction effect, comparing the expression ratio of the transcript over all other transcripts within a gene between conditions [11]. For the gene-level test, the transcript-level p-values are aggregated to gene-level q-values using the `perGeneQValue` function from DEXSeq [2], which amounts to controlling the FDR at level

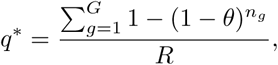

with G the number of genes, *n_g_* the number of transcripts for gene *g*, *θ* the significance threshold and *R* the number of rejections.

In the confirmation stage of the stage-wise analysis, we use the Shaffer MSRB method [36]. Genes in the confirmation stage have passed the screening stage, hence at least one of the transcripts should be differentially used between conditions. Since one transcript is differentially used, the difference in usage must be compensated for by at least one other transcript. Hence, all genes passing the screening stage should at least have two, and possibly more, DTU transcripts. According to the Sha er method the two most significant transcripts can be tested at a significance level of *α_II_*/(*n_g_* − 2) and from the third most significant transcript onwards the procedure reduces to the Holm method [37]. If a gene only consists of two transcripts, both are always called significant as soon as the gene passes the screening stage.

### Differential transcript expression

Isoform-level estimated counts are rounded to the closest larger integer and transcripts with no expression over all samples are discarded from the analysis. A negative binomial model is fit for every transcript using edgeR [23] and statistical inference is performed through likelihood ratio tests. For a transcript-level analysis the p-values are adjusted using BH correction while for a gene-level analysis they are aggregated to gene-level q-values as described in the previous section. Similar to the DTU analysis we account for the fact that genes in the confirmation stage must have at least one significant transcript, however there is no further dependency between the hypotheses for DTE. Therefore the Shaffer MSRB method only provides additional power for the most significant transcript, i.e. by testing it at Isoform-level estimated counts are rounded to the closest larger integer and transcripts with no expression over all samples are discarded from the analysis. A negative binomial model is fit for every transcript using edgeR [23] and statistical inference is performed through likelihood ratio tests. For a transcript-level analysis the p-values are adjusted using BH correction while for a gene-level analysis they are aggregated to gene-level q-values as described in the previous section. Similar to the DTU analysis we account for the fact that genes in the confirmation stage must have at least one significant transcript, however there is no further dependency between the hypotheses for DTE. Therefore the Sha er MSRB method only provides additional power for the most significant transcript, i.e. by testing it at *α_II_*/(*n* − 1) and from the second transcript onwards it reduces to the Holm [37] method.

### Case study

The unfilltered, unnormalized kallisto processed data was downloaded from The Lair project website (http://pachterlab.github.io/lair/) [46]. We rounded the kallisto estimated transcript counts to the closest larger integer and removed genes with only one transcript for DTU analysis and transcripts with no expression over all samples for both DTU and DTE analysis. differential transcript usage was assessed using DEXSeq [2] and DTE analysis was performed using edgeR [23]. A patient block effect was added to account for the correlation between control and tumoral tissue within patients and inference was performed as described in the simulation study. Both analyses were performed on a target 5% OFDR level. FWER correction in the confirmation stage of the stage-wise testing procedure was performed using an adapted Holm-Shaffer method [36], as described in the simulation study.

